# Capitalizing on Competition: An Evolutionary Model of Competitive Release in Metastatic Castration Resistant Prostate Cancer Treatment

**DOI:** 10.1101/190140

**Authors:** Jeffrey West, Yongqian Ma, Paul K. Newton

**Author notes:** corresponding author *Email addresses:* (Jeffrey West), (Yongqian Ma), (Paul K. Newton).

## Abstract

The development of chemotherapeutic resistance resulting in tumor relapse is largely the consequence of the mechanism of competitive release of pre-existing resistant tumor cells selected for regrowth after chemotherapeutic agents attack the previously dominant chemo-sensitive population. We introduce a prisoners dilemma mathematical model based on the replicator of three competing cell populations: healthy (cooperators), sensitive (defectors), and resistant (defectors) cells. The model is shown to recapitulate prostate-specific antigen measurement data from three clinical trials for metastatic castration-resistant prostate cancer patients treated with 1) prednisone, 2) mitoxantrone and prednisone and 3) docetaxel and prednisone. Continuous maximum tolerated dose schedules reduce the sensitive cell population, initially shrinking tumor volume, but subsequently “release” the resistant cells to re-populate and re-grow the tumor in a resistant form. Importantly, a model fit of prostate data shows the emergence of a positive fitness cost associated with a majority of patients for each drug, without predetermining a cost in the model a priori. While the specific mechanism associated with this cost may be very different for each of the drugs, a measurable fitness cost emerges in each. The evolutionary model allows us to quantify responses to conventional therapeutic strategies as well as to design adaptive strategies.

## 1. Introduction

In his now classic 1961 study of competition for space between two species of barnacles in the intertidal zone off the Scottish coast, Joseph Connell [1] discovered something interesting. The blue barnacles *Balanus* normally occupied the intertidal zone, while the brown barnacles *Chthamalus* occupied the coast above high tide. Despite the commonly held belief that each occupied their own niche because of different adaptations to local micro-conditions, Connell hypothesized that the colonization of the intertidal zone by *Balanus* was actually preventing *Chthamalus* from inhabiting this region. To test this, he removed the blue barnacles from the intertidal zone and tracked the subsequent penetration of *Chthamalus* into this region. He concluded that *Chthamalus* had undergone *relief from competition* with *Balanus* which allowed it to flourish where previously it could not. The point, he emphasized, was there was nothing *inherent* about the micro-environment of the intertidal zone that was preventing *Chthamalus* from occupying this region. It was simply the competition against a more dominant species that was holding it back. Without the presence of that species, *Chthamalus* happily claimed both zones as fundamental niches. Thus, the important notion of *competitive release* was formulated (see Grant [2]). When two (or more) sub-species compete for the same resources, with one species dominating the other, if the dominant species is removed, this can provide the needed release from competition that can allow the less dominant species to flourish. The mirror image of competitive release is the related notion of *character displacement* developed by Brown and Wilson [3] in which competition can serve to displace one or more morphological, ecological, behavioral, or physiological characteristics of two closely related species that develop in close proximity. These concepts are now well established as part of the overall framework of co-evolutionary ecology theory and play an important role in the evolution of chemotherapeutic resistance in cancer [4, 5, 6, 7].

Since co-evolution among competing subclones is now a well established [8] process in malignant tumors, the mechanism of competitive release should be expected to play a role and affect the chemotherapeutic strategies one might choose to eliminate or control tumor growth. Indeed, tumor relapse and the development of chemo-therapeutic resistance is now thought largely to be a consequence of the evolutionary mechanism of competitive release of pre-existing resistant cells in the tumor which are selected for growth after chemotherapeutic agents attack the subpopulation of chemo-sensitive cells which had previously dominated the collection of competing subclones. This concept, perhaps most forcefully advocated by Gatenby and collaborators [9], is gaining acceptance by clinicians. A recent (2012) systematic literature analysis of cancer relapse and therapeutic research showed that while evolutionary terms rarely appeared in papers studying therapeutic relapse before 1980 (< 1%), the language usage has steadily increased more recently, due to a huge potential benefit of studying therapeutic relapse from an evolutionary perspective [10]. Anticancer therapies strongly target sensitive cells in a tumor, selecting for resistance cell types and, if total eradication of all cancer cells is not accomplished, the tumor will recur as derived from resistant cells that survived initial therapy [11]. It is argued by Gatenby [9] that eradicating most disseminated cancers may be impossible, undermining the typical goal of cancer treatment of killing as many tumor cells as possible. The underlying assumption of this approach has been that a maximum cell-kill will either lead to a cure or, at worst, maximum life extension. Taking cues from agriculturists who have long abandoned the goal of complete eradication of pests in favor of applying insecticides only when infestation exceeds a threshold in the name of “control” over “cure,” there are those who advocate for a shift from the cure paradigm in cancer treatments to a control paradigm [9, 12].

### 1.1. The likelihood of pre-existing resistance

Pre-existing resistant sub-clones should generally be present in all patients with late-stage metastatic disease (for single point mutations which confer resistance), a conclusion supported by probabilistic models [13] and from tumor samples taken prior to treatment [14, 15] which have been reported for melanoma [16], prostate cancer [17], colorectal cancer [18, 19], ovarian cancer [20], and medulloblastoma [21]. According to this view, treatment failure would not be due to *evolution* of resistance due to therapy, but rather the pre-existing presence of resistant phenotypes that are relatively sheltered from the toxic effects of therapy [7].

The likelihood of pre-existing resistance has important therapeutic implications. If we assume no pre-existing resistance, then most models predict maximum dose-density therapy will reduce the probability of resistance largely because this treatment minimizes the number of cell-divisions, thereby minimizing the risk of a mutation leading to acquired resistance [7]. By contrast, in pre-existing resistance scenarios, the maximum dose-density therapy strategy lends itself to competitive release due to the evolutionary nature of tumor progression. Most pre-clinical efforts that aim to maximize the short-term effect of the drug on sensitive cells does not significantly affect the long-term control of cancer [13]. This is because the phenomenon of competitive release can occur via the harsh selective pressure imposed by the tumor microenvironment after cancer therapies diminish the presence of the dominant (i.e. the chemo-sensitive) clone. Additionally, the process of metastasis may allow a resistant subclone in the primary tumor to emerge elsewhere [22].

### 1.2. Leveraging the cost of pre-existing resistance

Pre-existing mutations that are responsible for conferring resistance may be associated with a phenotypic cost, or a reduced fitness, compared to the average fitness of the sensitive cell population [23]. Even factoring in this fitness cost, deleterious mutations are still expected to be present in late-stage metastatic cancers [24]. This cost can come in many ways, such as an increased rate of DNA repair, or an active pumping out of the toxic drug across cell membranes. All of these strategies use up a finite energy supply that would otherwise be available for invasion into non-cancerous tissues or proliferation. The rapid removal of chemo-sensitive cells during therapy releases the resistant population from unwanted competition and thereby permits unopposed proliferation of the resistant cell population. In contrast, the goal of an adaptive therapy is to maintain a stable tumor burden that permits a significant population of chemo-sensitive cells for the purpose of suppressing the less fit but chemo-resistant populations, consistent with the philosophy that it takes an evolutionary strategy to combat an evolving tumor [22].

A theoretical framework for these adaptive therapies first developed by Gatenby [23], leverages the notion that pre-existing resistance is typically present only in small population numbers due to a cost of resistance. This less fit phenotype is suppressed in the Darwinian environment of the *untreated* tumor but treatments that are designed to kill maximum numbers of cells remove the competition for the resistant population and ultimately select for that population during tumor relapse^1^. While the goal of an adaptive therapy (to capitalize on the cost of resistance by maintaining a stable sensitive cell population in order to suppress the resistant population) has gained some level of acceptance, the ideal adaptive therapy schedule in practice, for different tumor types and growth rates is far from settled. Gatenby’s paper modulates dose density only, while stating that an ideal adaptive therapy would also modulate the drug, as well as the dose schedule (both dose and density) [23]. Some of these evolutionary ideas were tested experimentally using mouse models with modulated dose strength and dose vacations designed to maintain a stable, controllable tumor volume [25, 26]. This two-phase adaptive therapy involved an initial high-dose phase to treat the exponential growth of the tumor and a second phase designed for stable tumor control using a variety of strategies (such as decreasing doses or skipping doses when stability is achieved). Several spatial, agent-based computational models have modulated dose strength with respect to a threshold value of tumor size (a fraction of the original tumor burden) [27, 28]. Findings suggest that adaptive therapies based on evolutionary treatment strategies that maintain a residual population of chemo-sensitive cells may be clinically viable, and is currently extended to an on-going clinical trial (NCT02415621) which adaptively controls the on/off cycling of abiraterone [29].

With these advances in mind, the goal of this manuscript is to use the evolutionary framework introduced and advocated over the past 10 years [23, 9, 27, 28, 29] to mathematically model the important parameters of competitive release and use that framework to better understand therapeutic implications of the cost of developing resistance and to learn how to exploit this cost that the tumor pays. Specifically, we introduce a three-component replicator system with a prisoner’s dilemma payoff matrix [30] to model the three relevant subclonal populations: healthy cells (H), sensitive cells (S), and resistant cells (R). Using the nullcline information in a triangular phase plane representation of the nonlinear dynamics of the system, we first show the essential ingredients that render competitive release possible. Then, using the parameters that control selection pressure (hence relative growth rates) on the three subclonal populations, we attempt to maintain the tumor volume at low levels so that the resistant population does not reach fixation.

### 1.3. Retrospective analysis of metastatic castration-resistant prostate cancer

A recent retrospective analysis of tumor measurement data (PSA levels) from eight randomized clinical trials with metastatic castration-resistant prostate cancer (mCRPC) used a simple linear combination of exponentials model to estimate the growth and regression rates of disease burden over time [31]. In total, over 67% of patients were fit to models with a positive regrowth rate, indicating failure due to resistance. Prostate-specific antigen (PSA) measurement data for patients in each treatment silo (prednisone only, mitoxantrone and prednisone, docetaxel and prednisone) were obtained through the Project Data Sphere open data portal (https://www.projectdatasphere.org), and we show that this model is able to adequately fit data for each treatment type with the additional capability of allowing us to track responses to conventional therapeutic strategies and design new adaptive strategies as the tumor evolves. The model can be used to test previously proposed adaptive therapies, but we propose a novel schedule utilizing quantitative tools from nonlinear dynamical systems theory which use the current global state of the nonlinear replicator system with respect to the nullcline curves of the equations as well as parameters controlling relative fitness levels of the competing sub-populations. The simulated chemotherapeutic strategies that we implement, based on tracking the phase-space structure of of replicator system, are ones that can adapt on the same timescale as the inherent timescale of evolution of the subclones comprising the tumor, i.e. are as dynamic as the tumor. While this approach cannot be preplanned by the oncologist at the beginning of therapy like classical strategies, we provide discussion to explain how an evolutionary game theory model describing the fitness landscape (described below) is useful to understand the underlying features of a dynamical fitness landscape associated with a cost to resistance: a three-way prisoner’s dilemma. Specifically, the model indicates a boundary over which an adaptive therapy will cease to be effective. Our model focuses on cost to resistance, as opposed to specific mechanisms of resistance (i.e. hormone resistance, for example). This has the advantage in some ways as being agnostic to resistance mechanisms, but on the other hand, in this manuscript, we can not distinguish between different mechanisms: the analysis explores the effect of a fitness cost on the competitive release phenomenon. Importantly, this analysis provides insight into the resistance cost of three common therapies used in mCRPC, as we show that a fit of our model to prostate data showed the *emergence* of a positive cost value (described below) in a majority of patients, without predetermining a cost in the model *a priori*. We now describe the model.

## 2. Materials and Methods

A schematic of a finite-population three component model (healthy, sensitive, resistant) of competitive release is shown in figure 1, where the tumor consisting of sensitive and resistant cells is competing with the surrounding healthy tissue. At diagnosis (see figure 1, left), the tumor is dominated by sensitive cells (red) which out compete the surrounding healthy population (blue) during unhindered tumor progression. A small portion of resistant cells (green) remains in small numbers, suppressed by the larger sensitive population. After several rounds of chemotherapy, the tumor shrinks, leaving the resistant population largely unaffected (figure 1, middle). Inevitably, the tumor relapses due to the small number of sensitive cancer cells remaining after therapy (figure 1, right). In the absence of competition from the dominant sensitive population, the resistant cells grow unhindered, rendering subsequent rounds of chemotherapy less effective. Subsequent application of identical therapies will have a diminished effect. Figure 2 shows the process in a ‘Muller fishplot’, which we will use later to track the subclonal populations. This representation was first utilized in cancer to compare modes of clonal evolution in acute myeloid leukemia (see [32]) and is particularly useful in computational models where every cell type can be tracked in time. A fishplot shows the tumor burden (vertical axis) over time (horizontal axis) and the clonal lineages (a subclone is encased inside of the founding parent clone in the graph). The schematic depicts unhindered tumor growth after the first driver mutation (figure 2, left) where the tumor grows exponentially before diagnosis, during which time a resistant mutation occurs (figure 2, middle). After diagnosis (dashed line), a regimen of continuous chemotherapy shows initial good response and tumor regression, but the resistant population grows back (although at a slower growth rate) unhindered by competition, leading to relapse (figure 2, right). Previously, a linear combination of exponentials model has been proposed to track the relative tumor volume, *v*(*t*), after treatment as a function of the exponential death rate of the sensitive cells, *d*, the exponential growth rate of the resistant cells, *g*, and the initial fraction of resistant cells, *f* [13]. The model can be written as follows:

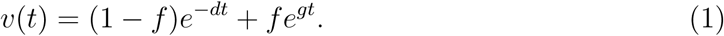

This model, shown to be a reasonably good description of the changing tumor size during therapy for colorectal, prostate, and multiple myeloma cancers, identifies the important parameters in competitive release: initial fractional resistance (*f*), and birth/death rates (*g*,*d*) for the resistant and sensitive populations, respectively. The model is used to fit prostate-specific antigen (PSA) measurement data from retrospective analysis of three randomised clinical trials with metastatic castration-resistant prostate cancer to estimate the growth (g) and regression (d) rates of disease burden over time. Four representative patients are chosen from the control arms of each randomized trial and shown in figure 3: treatment with prednisone only [33] (left column: figures 3a,3d,3g,3j); treatment with mitoxantrone and prednisone [34] (middle column: figures 3b,3e,3h,3k) and treatment with docetaxel and prednisone [35] (right column: figures 3b,3e,3h,3k). PSA data and model fits are normalized by *v*(*t* = 0) (black dots) and exponential fits are shown in blue.

**Figure 1:**
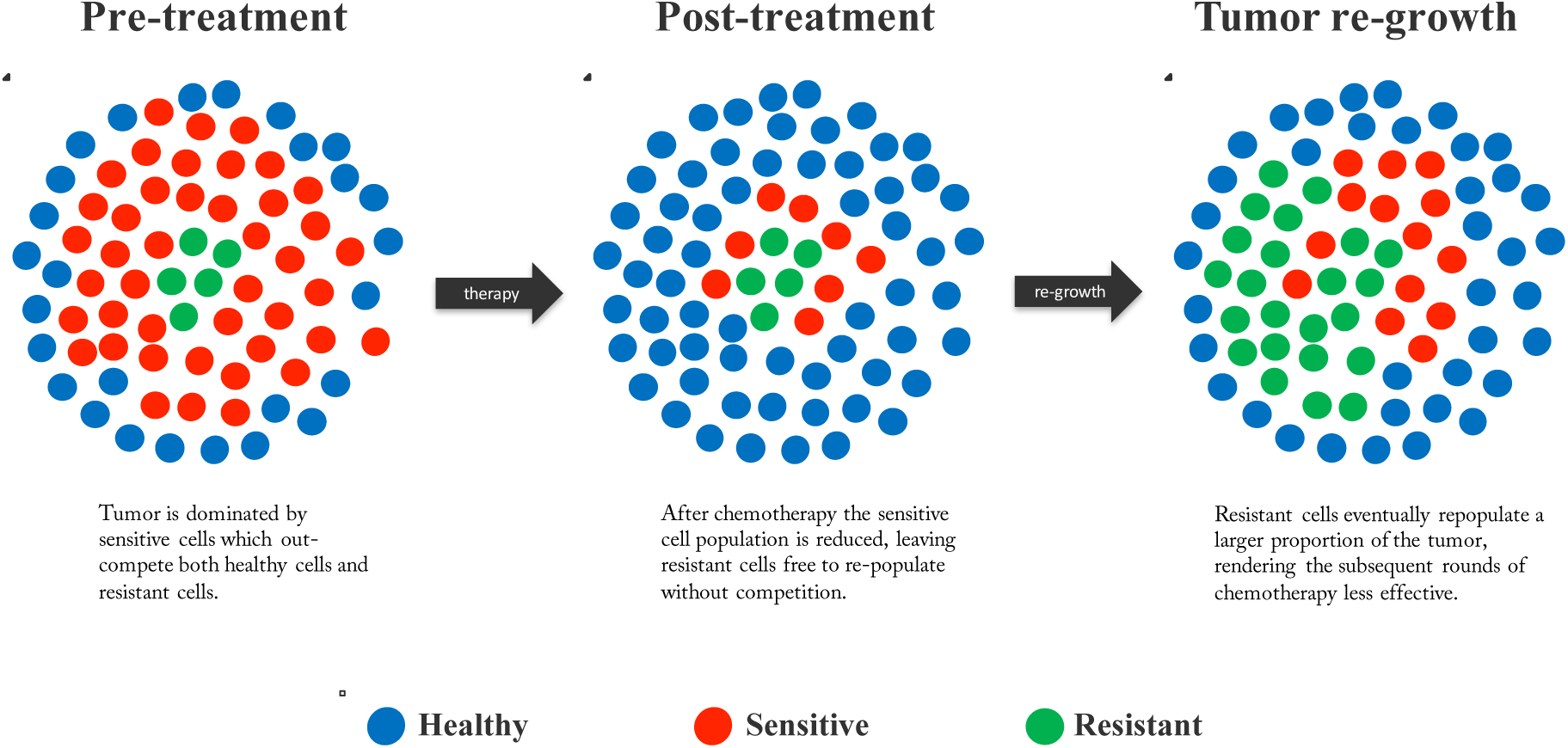
Schematic of competitive release in a tumor. — (a) Prior to treatment, a tumor consists of a large population of sensitive cells (red) and a small population of less fit resistant cells (green) competing for resources with the surrounding healthy cells (blue); (b) Chemotherapy targets the sensitive population (middle), selecting for the less fit resistant population that thrives in the absence of competition from the sensitive population; (c) Upon regrowth, the tumor composition has larger numbers of resistant cells, rendering the subsequent rounds of treatment less effective.

**Figure 2:**
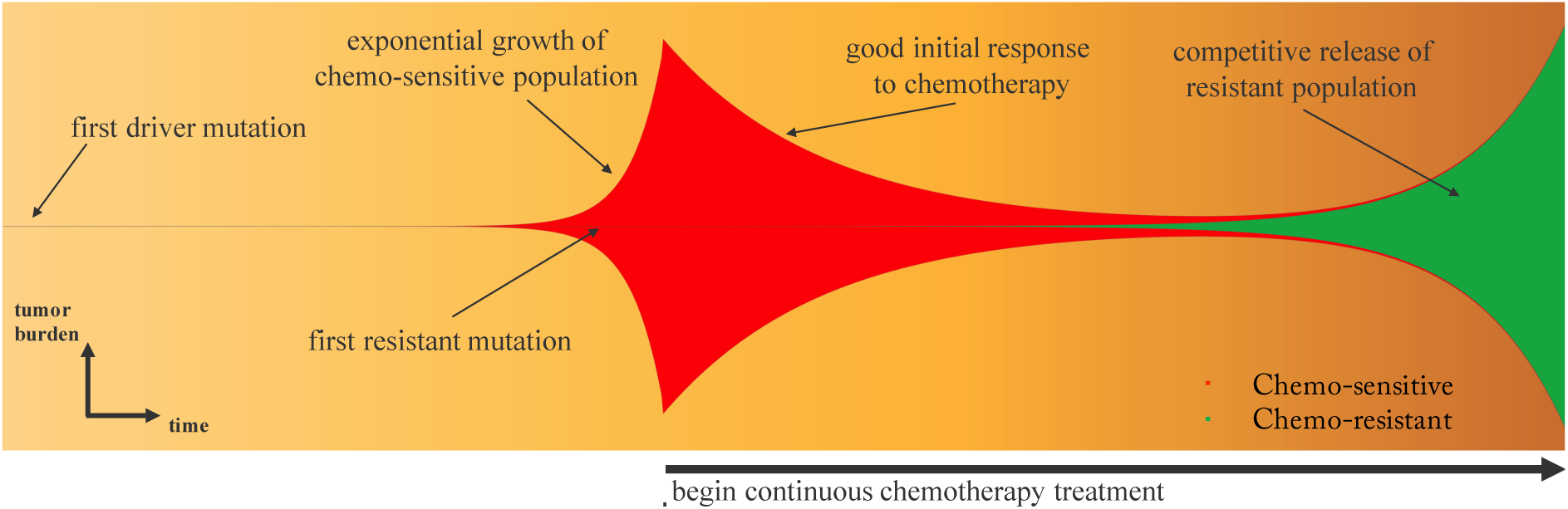
Clonal evolution of competitive release. — A fishplot (sometimes known as a Müller plot), showing the tumor size (vertical axis) and composition (sensitive: red; resistant: green) over time (horizontal axis, left to right) with important events annotated. After first driver mutation (left), initial exponential growth of sensitive population occurs until diagnosis (dashed line). Continous therapy targeting the chemo-sensitive population responds well with a decrease in tumor burden. In the absence of sensitive cells, the resistant population (existing in small numbers before the start of therapy) grows to become the dominant clone at relapse, albeit typically with lower exponential growth rate.

**Figure 3:**
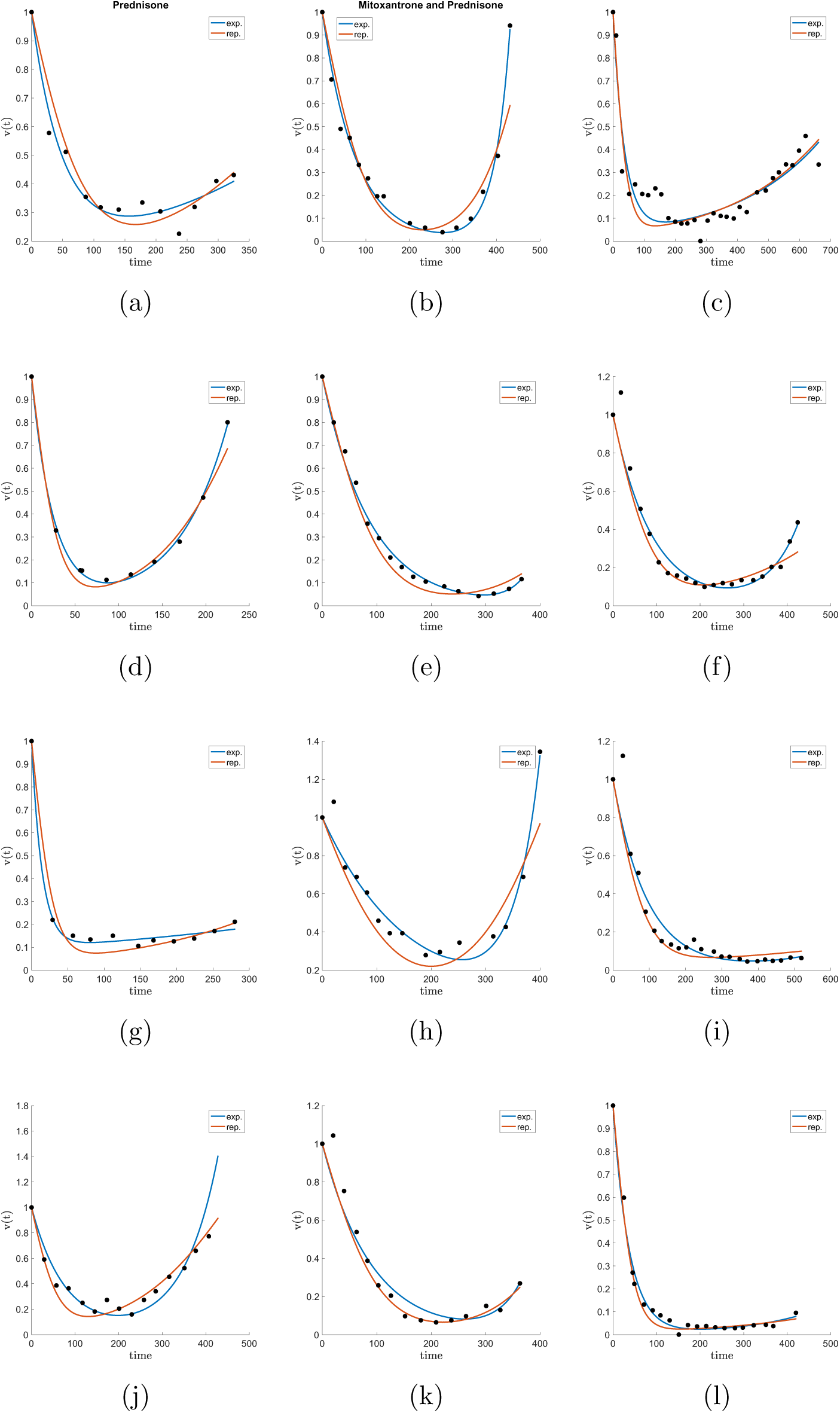
Model fits of PSA measurement data for metastatic castration-resistant prostate cancer. — Prostate-specific antigen (PSA) measurement data of four representative patients from three randomised clinical trials with metastatic castration-resistant prostate cancer. Left column: treatment with prednisone only [33]; middle column: treatment with mitoxantrone and prednisone [34]; right column: treatment with docetaxel and prednisone [35]. PSA data is normalized by *v*(*t* = 0) (black dots). The data is fit using the exponential model (eqn. 1; blue curve) and the replicator model (eqns 2, 3). Each patient is fit reasonably well with both models. Data was fit by parameter sweep of cost (*α* – *β*, eqn. 14, 15), initial fractional resistance *f* and selection pressure *w*. Parameters used are *α* – *β* = [0.02, 0.07, 0.20, 0.00, 0.20, 0.00, 0.20, 0.03, 0.20, 0.01, 0.15, 0.19]; *f* = [0.20, 0.02, 0.09, 0.05, 0.03, 0.07, 0.10, 0.10, 0.09, 0.10, 0.03, 0.03]; *w* = [0.10, 0.20, 0.20, 0.30, 0.20, 0.10, 0.30, 0.15, 0.10, 0.15, 0.20, 0.20] for a - l respectively.

Despite the fact that the model (1) curve-fits data reasonably well (labeled “exp.” in figure 3), it contains no evolutionary information or concepts, a keystone principle behind competitive release. We also include in the figure 3 fits of our model presented here (eqns. 2, 3) in red (labeled “rep.” in figure 3). Our evolutionary model is able to capture similar trends as the exponential model of equations (eqn. 1) but has the important property of allowing us to calculate the fitness cost associated with model fit. While each drug (columns) may have very different resistance *mechanisms*, ten of the twelve patients shown have a positive cost (*α* – *β*, described below). In the last section we will describe the dynamical phase portrait associated with a cost to resistance and implement adaptive strategies to capitalize on this cost that the tumor pays in order to maintain and grow its resistant population.

### 2.1. The replicator equation model

The dynamics of the fitness landscape of three competing cell types are described by the replicator equation (see [36]), which is a deterministic birth-death process in which birth and death rates are functions of cell fitness, and cell fitness is a function of prevalence in the population. Each *i*th cell type (*i* = 1, 2, 3) competes according to equation 2, where *x*_1_, *x*_2_, *x*_3_ are the corresponding frequency of healthy (H), sensitive (S) and resistant (R) cells, respectively, such that Σ_*i*_ *x*_*i*_ = 1.

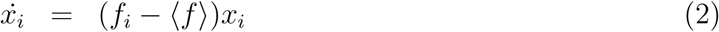

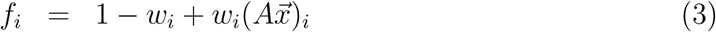

Here, 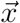 is the vector 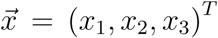 and (*Ax*)_*i*_ is the *i*th element of vector *Ax*. The prevalence of each sub-population, *x*_*i*_, changes over time according to the changing population fitness, *f*_*i*_, as compared to the average fitness of all three populations 〈*f*〉 = *f*_1_*x*_1_ + *f*_2_*x*_2_ + *f*_3_*x*_3_. If the fitness of the sub-population is greater than the average fitness (*f*_*i*_ – 〈*f*〉 > 0), that sub-population grows exponentially, whereas if it is less (*f*_*i*_ – 〈*f*〉 < 0), it decays.

Before therapy, each subpopulation (healthy, chemo-sensitive, and resistant cells) the selection pressure is constant across all cell types (i.e. *w*_*i*_ ≡ *w*, *i* = 1, 2, 3) at a level that represents the natural selection pressure the tumor environment imposes on the different subpopulations. These values discussed in the literature are typically small, in the range *w*_*i*_ ≡ *w* ≈ 0.1 – 0.3. We implement chemotherapy in our model by changing the selection pressure parameters on each of the subpopulations of cells. Therapy can be administered at different doses (i.e. values of the drug concentration: *c*; 0 ≤ *c* ≤ 1). A higher value of c indicates a stronger dose of chemotherapy drug (described in more detail in [37]). This follows the schematic in figure 4 which depicts the change in the fitness landscape before and after therapy. In figure 3, dose concentration is assumed to be constant for a specific drug while patient-specific parameters are the selection pressure (*w*), cost of resistance (discussed below), and initial fraction of resistant cells (*f*). Values are altered as follows (see figure 4 for explanation of changing fitness landscape):

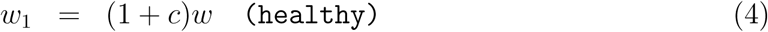

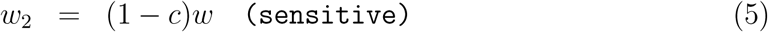

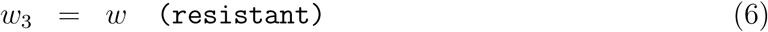

The fitness landscape (eqn. 3) is described in detail by the entries of the payoff matrix *A*, (eqn. 7) where each pairwise cell-cell interaction is described by the row and column values, which are parameters in the fitness equation (3). The equation can be separated into three pairwise games: (*H*,*S*), (*H*,*R*), or (*R*, *S*), which are all calculated using a prisoner’s dilemma (cooperators, defectors) game. This necessitates the following inequalities of the payoff matrix below (eqn. 7): *h* > *a* > *j* > *b*, *l* > *a* > *n* > *o*, and *k* > *n* > *j* > *m*. With this paradigm, the cancer cells (sensitive or resistant) act as ‘defectors’, whereas the healthy cells act as ‘cooperators’ in each interaction. More discussion of why the prisoner’s dilemma matrix, which models the evolution of defection, is a useful paradigm for cancer can be found in [30, 38].

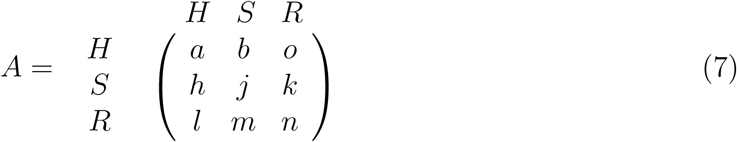

### 2.2. The linearized system and the cost of resistance

The notion of the *cost of resistance* is highlighted in figure 4. With no therapy, the sensitive cells exhibit fastest growth due to their higher fitness value relative to both the resistant population and the healthy population. The difference between the baseline fitness values of the sensitive cells and the resistant cells can be thought of as the ‘price paid’ by the resistant population to retain their resistance to toxins. This cost, in our model, is quantified as the difference in the (linearized) growth rates of the two populations (type 2: sensitive; type 3: resistant). Linearizing eqn (2), (3), (which form a cubic nonlinear system if expanded out) gives rise to the sensitive-resistant uncoupled system:

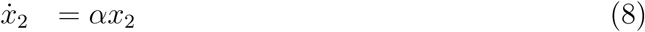

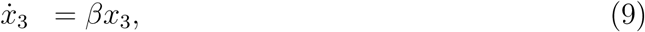

with the growth parameters:

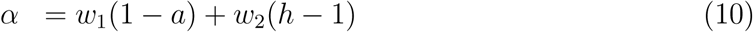

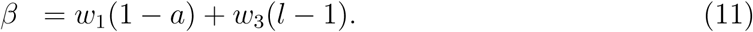

Using eqns (4), (5), (6) gives:

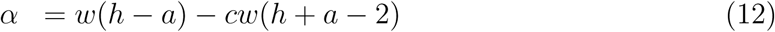

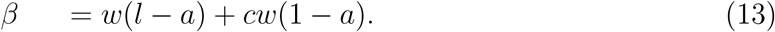

With no therapy, *c* = 0, we have:

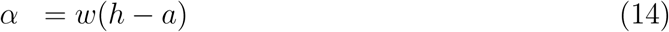

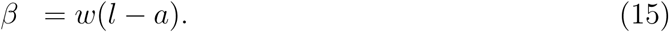

We call the fitness cost of resistance the difference between these growth rates with no therapy, hence (*α* – *β*) = *w*(*h* – *l*).

**Figure 4:**
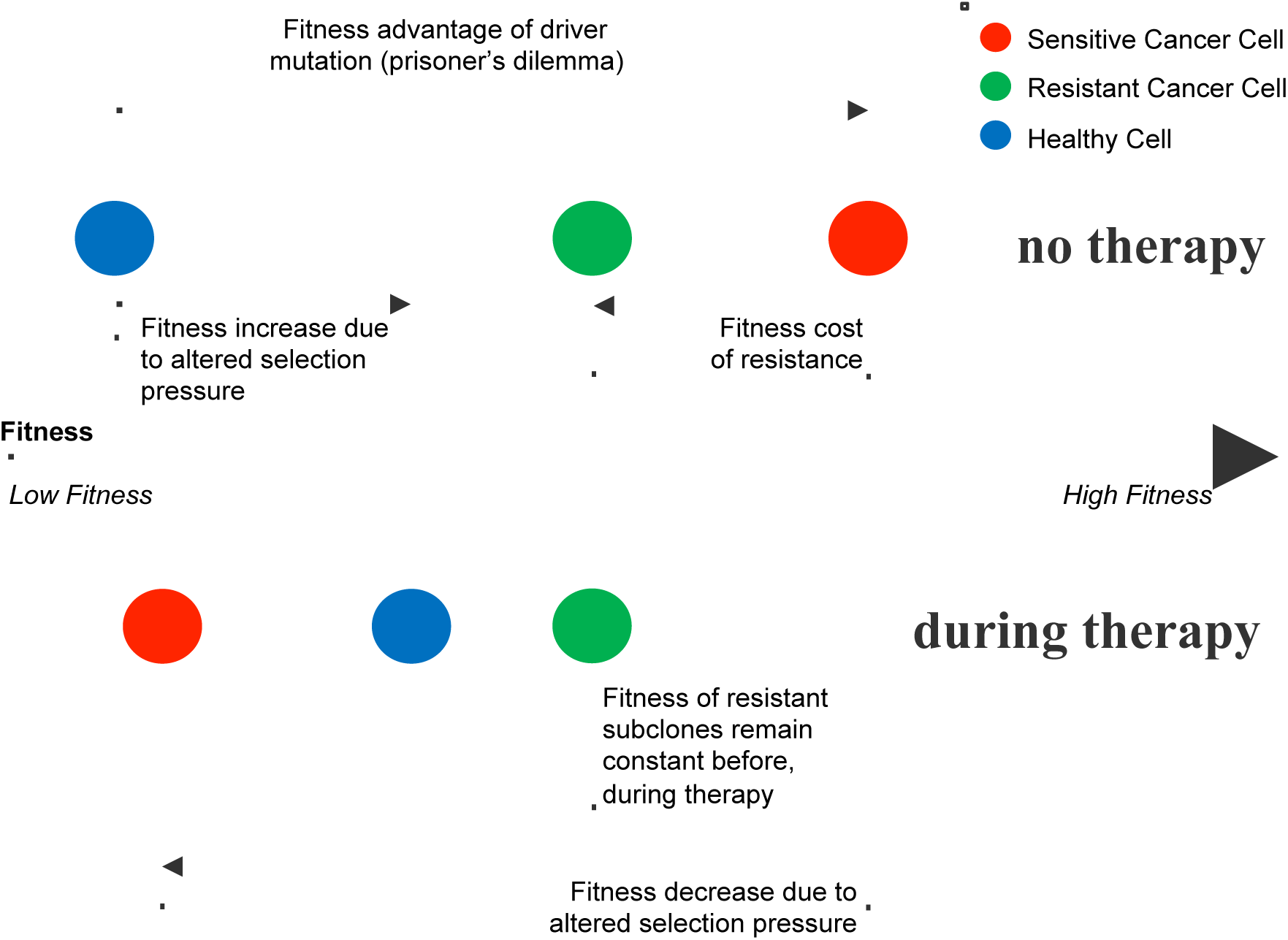
Fitness landscape before and during therapy. — A schematic of the fitness of each subpopulation before therapy (top) and during therapy (bottom). A driver mutation leads to a fitness advantage of the cancer cell (red), determined by the prisoner’s dilemma payoff matrix. A subsequent resistant-conferring mutation comes at a fitness cost (green). The fitness of the resistant population is unaffected by therapy’s selective pressure, but the healthy population is given an advantage over the chemo-sensitive population.

## 3. Results

It is useful to view the nonlinear dynamical trajectories of the system using the tri-linear coordinates shown in figure 5a, which gives a representation of the clonal phase space for every possible value of 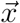 [39]. The corners represent saturation of a single cell type (e.g. the top corner represents 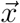 = [1, 0, 0], or all healthy cells. Figure 5b shows the nullcline information of the dynamical system (curves for which *x→* = 0) with therapy off (solid green line) and on (dashed green line). As the trajectory crosses a particular nullcline, the growth (*ẋ*_*i*_ > 0) / decay (*ẋ*_*i*_ < 0) on one side switches to decay/growth, allowing for the possibility of trapping an orbit in a closed loop for a finite period of time if the state of the system can be ‘steered’ appropriately. We do this by altering the dose concentration parameter *c* (a parameter that can be accessed clinically) in eqns (4) in an off-on (bang-bang) fashion, (eqn. 5, 6). This is schematically depicted in figure 5b. The dynamics of equation (2) are shown in state space diagrams in Figure 5 for no therapy (figure 5c: *c* = 0) and with therapy (figure 5d: *c* = 0.6). Before treatment, the healthy (H; top corner), sensitive (S; bottom left corner), and resistant (R; bottom right corner) populations compete according to equation (2) and follow trajectories shown (black) in figure 5c. Instantaneous relative velocity is indicated by background color gradient (red to blue). All internal trajectories (pre-therapy) lead to tumor growth and eventual saturation of the sensitive population (bottom left corner). The resistant population nullcline (line of zero growth; *ẋ*_*R*_ = 0) is plotted in dashed dark red in figure 5c. With no therapy (left), the nullclines divide the triangle into 3 regions.

- Region 1: *ẋ*_*H*_ > 0 *ẋ*_*S*_ > 0 *ẋ*_*R*_ < 0;
- Region 2: *ẋ*_*H*_ < 0 *ẋ*_*S*_ > 0 *ẋ*_*R*_ < 0;
- Region 3: *ẋ*_*H*_ < 0 *ẋ*_*S*_ > 0 *ẋ*_*R*_ > 0.

**Figure 5:**
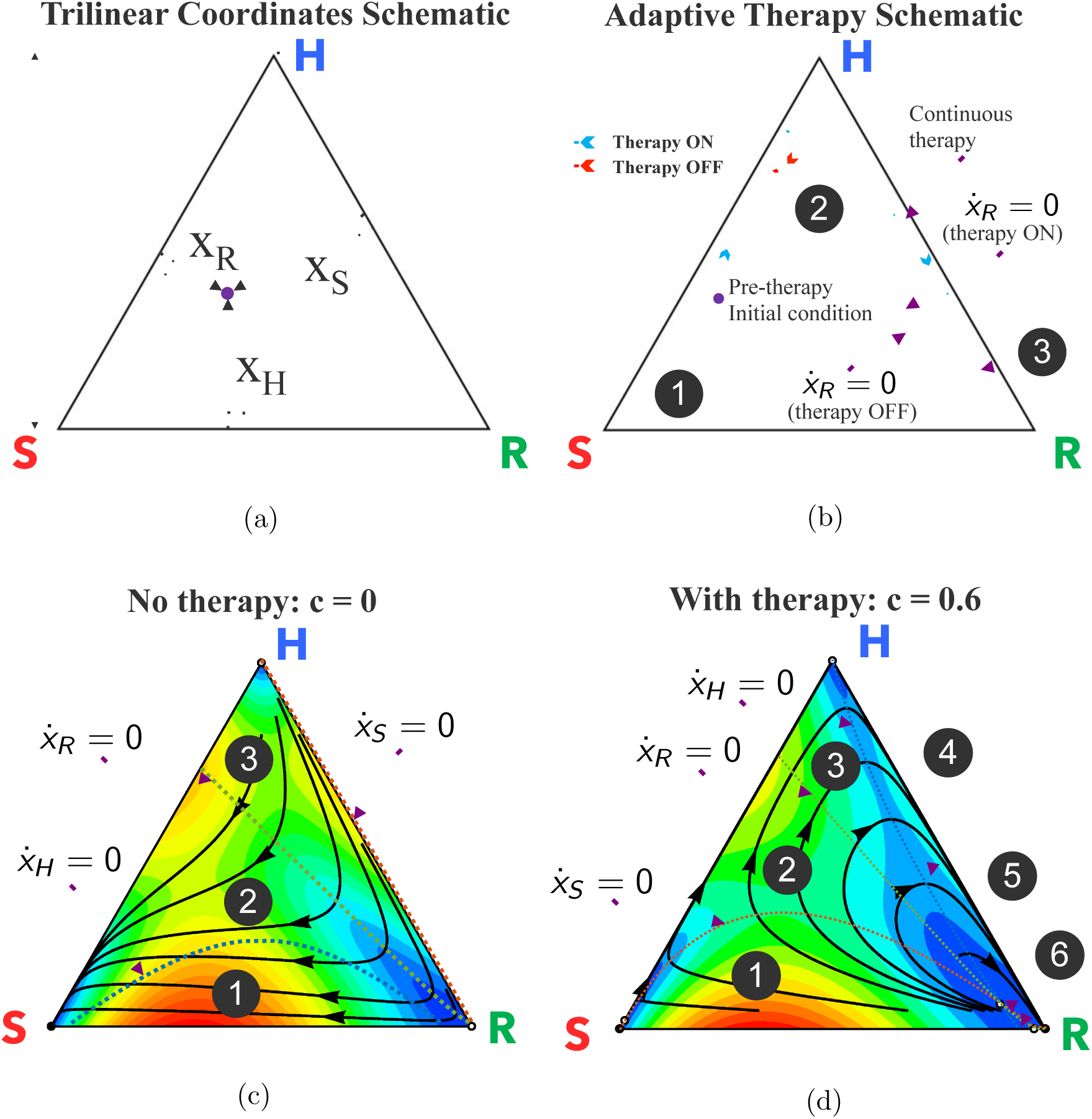
Dynamic phase portraits before and during chemotherapy. — (a) Trilinear coordinate phase space representation; (b) Schematic of proposed adaptive therapy concept using the resistant nullclines to determine therapy “on” and “off’ times in order to trap the tumor in the controllable region 2, and reach approximate cycle that repeats back on itself in red. The continuous therapy is also plotted in dashed blue, for comparison. Two nullclines divide the triangle into 3 regions; region 1: *ẋ*_*R*_ < 0 for both therapy on and off; region 2:*ẋ*_*R*_ > 0 for therapy off and *ẋ*_*R*_ < 0 for therapy on; region 3: *ẋ*_*R*_ > 0 for both therapy on and off. (c) Before chemotherapy, the healthy (H), sensitive (S), and resistant (R) populations compete on a dynamical fitness landscape, with several solution trajectories shown (black) and the instantaneous relative velocity indicated by background color gradient (red to blue). All internal trajectories lead to tumor growth and eventual saturation of the sensitive population (bottom left corner). Each population nullcline (line of zero growth: *ẋ*_*i*_ 0) is plotted: healthy (dashed blue), sensitive (dashed red), and resistant (dashed green). The nullclines divide the triangle into 3 regions. Region 1: *ẋ*_*H*_ > 0 *ẋ*_*S*_ > 0 *ẋ*_*R*_ < 0; Region 2: *ẋ*_*H*_ < 0 *ẋ*_*S*_ > 0 *ẋ*_*R*_ < 0; Region 3: *ẋ*_*H*_ < 0 *ẋ*_*S*_ > 0 *ẋ*_*R*_ > 0; (d) Chemotherapy alters the selection pressure to the disadvantage of chemo-sensitive cancer population and advantage of the healthy population (shown for *c* = 0.6, *α* = 0.020, *β* = 0.018, *w* = 0.1). In this case, the nullclines divide the triangle into 6 regions; Region 1: Region 1: *ẋ*_*H*_ > 0 *ẋ*_*S*_ > 0 *ẋ*_*R*_ < 0; Region 2: *ẋ*_*H*_ > 0 *ẋ*_*S*_ < 0 *ẋ*_*R*_ < 0; Region 3: *ẋ*_*H*_ > 0 *ẋ*_*S*_ < 0 *ẋ*_*R*_ > 0;Region 4: *ẋ*_*H*_ < 0 *ẋ*_*S*_ < 0 *ẋ*_*R*_ > 0; Region 5: *ẋ*_*H*_ < 0 *ẋ*_*S*_ > 0 *ẋ*_*R*_ > 0; Region 6: *ẋ*_*H*_ < 0 *ẋ*_*S*_ > 0 *ẋ*_*R*_ < 0; Solution trajectories (black) show initial trajectory toward healthy saturation (triangle top) but eventual relapse toward resistant population (bottom right of triangle) upon passing the resistant nullcline.

With chemotherapy (right) the selection pressure is altered to the disadvantage of chemo-sensitive cancer population and advantage of the healthy population (shown for *c* = 0.6, *α* = 0.020, *β* = 0.018, *w* = 0.1). In this case the nullclines divide the triangle into 6 regions.

- Region 1: *ẋ*_*H*_ > 0 *ẋ*_*S*_ > 0 *ẋ*_*R*_ < 0;
- Region 2: *ẋ*_*H*_ > 0 *ẋ*_*S*_ < 0 *ẋ*_*R*_ < 0;
- Region 3: *ẋ*_*H*_ > 0 *ẋ*_*S*_ < 0 *ẋ*_*R*_ > 0;
- Region 4: *ẋ*_*H*_ < 0 *ẋ*_*S*_ < 0 *ẋ*_*R*_ > 0;
- Region 5: *ẋ*_*H*_ < 0 *ẋ*_*S*_ > 0 *ẋ*_*R*_ > 0;
- Region 6: *ẋ*_*H*_ < 0 *ẋ*_*S*_ > 0 *ẋ*_*R*_ < 0.

Solution trajectories (black) show the initial trajectory toward healthy saturation (triangle top) but eventual relapse toward resistant population (bottom right of triangle) upon passing the resistant nullcline. The nullclines will be used later to determine timing schedules of adaptive therapy (see figure 6a).

**Figure 6:**
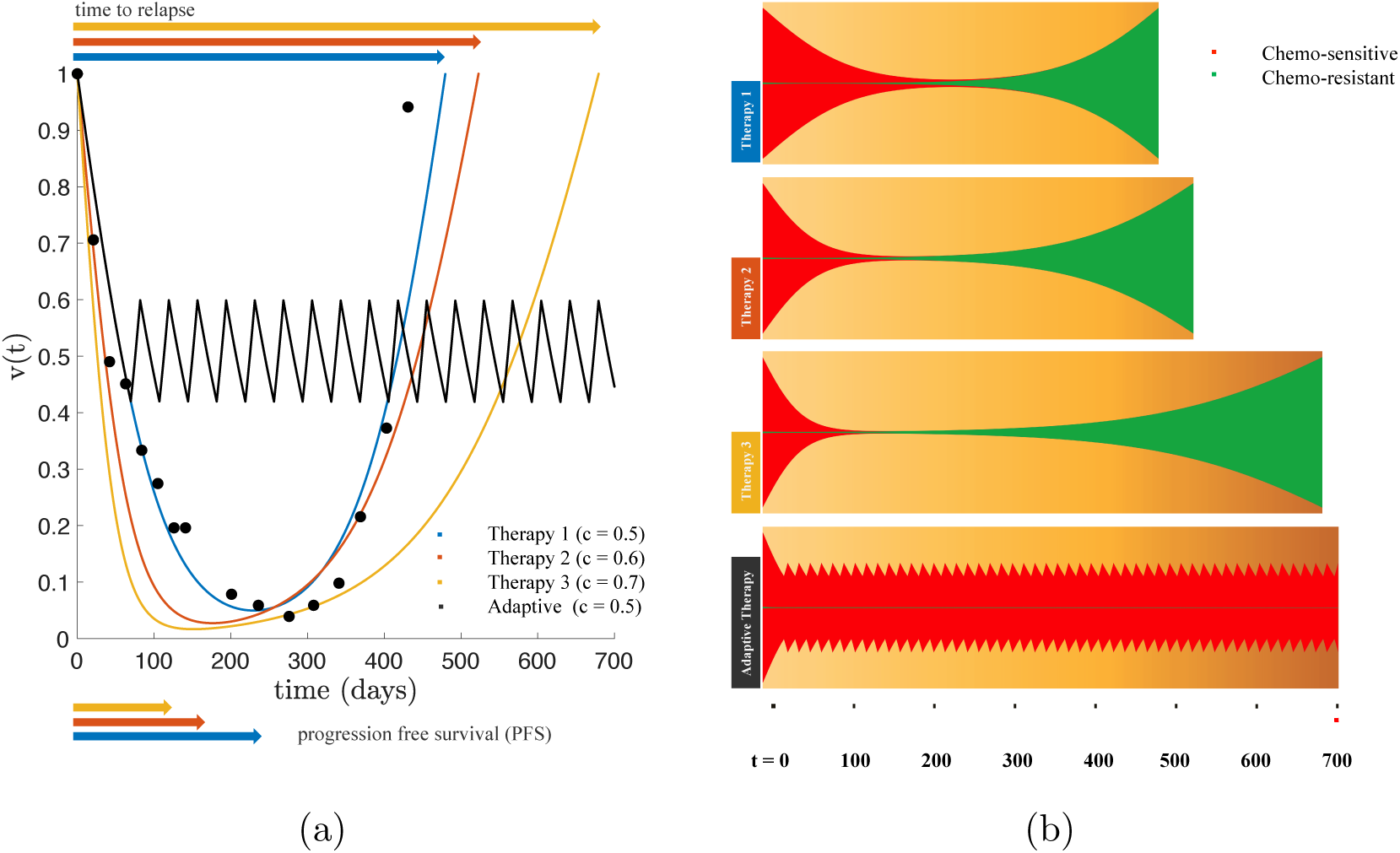
The effect of dose on tumor relapse and progression free survival under continuous and adaptive therapy. — (a) PSA data is from a single patient (see figure 3b) under continuous treatment (Mitoxantrone and Prednisone) is replotted (black dots) along with the best model fit (eqns. 2, 3) in blue. Using identical patient parameters, continuous treatment of two “new” drugs with higher effectiveness (i.e. increased effective dose, *c*) is shown in red and yellow. Time to relapse significantly increases with increasing dose while the progression free survival shows marginal, but decreasing, difference. An adaptive therapy (see figure 5b) is also simulated (solid black line), showing an increased control over the tumor; (b) The same four therapies are shown in a fish plot. Continous therapies show relapse to initial tumor size is dominated by chemo-resistant population (green). The adaptive therapy successfully suppressed the growth of the resistant population (bottom).

### 3.1. Managing competitive release

Figure 6 details the relationship between dose and two important measures of therapy effectiveness: progression free survival (PFS) and time to relapse. Measuring the effectiveness of a chemotherapy schedule based on the killing rate or progression free survival alone are not sufficient predictive measures of long-term cancer control [13]. PSA data from a patient who relapsed due to treatment resistance (figure 3b) is replotted in figure 6a (black dots) along with the model best fit of continuous treatment (blue; therapy 1). Also shown are new drugs with increased effectiveness of killing sensitive cells (red, yellow) simulated by increasing the effective dose concentration with identical patient-specific parameters (from the patient in figure 3b). Each increased dose corresponds to a slightly shorter PFS, but an increased time to relapse to the initial tumor volume. However, despite the increase in relapse times, none of these doses optimizes tumor control, as seen in the fishplots (figure 6, right). At the point of relapse to the initial tumor volume, the tumor is dominated by the presence of resistant clones (green), rendering future treatments ineffective. Oftentimes, the effectiveness of a new chemotherapy drug is be determined by PFS times when drugs that have high killing rates of sensitive cells may have *shorter* times to progression and lower total tumor burden at all times (everything else equal). The figure clearly shows that all treatments have similar progression free times but with a greater range of relapse times (even though continuous treatment always eventually leads to relapse).

An important detail emerges from the model: during treatment, the resistant nullcline is reached before the healthy nullcline (figure 5d). In other words, if the desire is to maintain treatment until the point of positive progression, this occurs when the trajectory is far past the resistant nullcline. In other words, appearances (based on tumor volume) can be deceiving - while the tumor may appear to be responding, the overall state may be well past the point of no return, and the resistant population is only preparing to re-populate. While all details of the ‘tumor phase space’ may not yet be directly measurable in the clinic, we propose that all successful adaptive therapies will operate in regions 1 and 2 in figure 5d; otherwise they ultimately will not be successful. In this way, meaningful insight is gained into the dynamics behind the cost to resistance, regardless of the mechanism of that resistance. Previous adaptive therapy schedules (described in the introduction; see [27, 28, 25, 29]) have benefited from not crossing this “hidden” resistant nullcline. We now propose one example of an adaptive therapy schedule which actively captures and uses information from the dynamic phase space.

A simple control paradigm is proposed to *indirectly* control the resistant population. Therapy targets only the chemo-sensitive cells, but the resistant population can be controlled by systematically choosing when to administer therapy and when to give drug holidays. Therapy “on” is for the purpose of killing sensitive cells. Therapy “off” is for the purpose of allowing a sufficient number of sensitive cells to remain, in order to suppress the resistant population. The control paradigm is as follows: a continuous dose of therapy is administered until the nullcline (*ẋ*_*R*_ = 0) is reached (see figure 5d, green dashed line). This is the starting point of positive growth for the resistant population (further therapy would result in *ẋ*_*R*_ > 0). At this point, a drug holiday (no therapy administered) is imposed until the second nullcline is reached (see figure 5c, green dashed line). The sensitive population is allowed to regrow until it is large enough to suppress the resistant population once again (and when *ẋ*_*R*_ = 0). Therapy is administered to allow the tumor to cycle back and forth between the two nullclines. This bang-bang (on-off) strategy allows an extension of relapse times. We emphasize that the specific times we turn the therapy on and off in this bang-bang strategy cannot be pre-planned, but depend on the position of the trajectory in the tumor phase space as the disease evolves.

This control paradigm is seen in figure 6a (solid black line) for identical initial conditions and identical drug dose. Rather than administer a continous dose, treatment holidays are given to trap the tumor trajectory between the two nullclines. As seen in the fishplot (figure 6b, bottom), the resistant population (green) is suppressed during the “off” times of drug holidays, leading to an extended time without relapse. This adaptive technique is successful for two reasons. First, the drug holidays allow an adequate sensitive population size to suppress the growth of the lower-fitness resistant population. Second, the resistant population is never allowed to reach a positive growth under treatment (*ẋ*_*R*_ > 0), therefore cannot take over the tumor cell population.

## 4. Discussion

The chemotherapeutic scheduling strategies outlined in this paper cannot be preplanned by the oncologist at the beginning of therapy like classical strategies [40], as they rely on significant decision making and continuous monitoring of the different subpopulations of cells that co-evolve as the tumor progresses. This means that the quality of the cell population monitoring system is crucial to the entire strategy, as has been pointed out in [41]. There can be no adaptive tumor control strategy without continuous monitoring of the sub-clones as it is not the tumor volume that is of primary interest, but the heterogeneous balance of the sub-clones comprising the tumor. In addition, the information gleaned from a detailed monitoring system cannot be acted upon unless the various administered drugs are sufficiently targeted to act efficiently and exclusively on specific sub-clones. These two systems must be in place (sensing and actuating) in order to successfully shape the fitness landscape and steer a growth trajectory in a desired direction. We also want to emphasize a separate point, which is that it is not enough to know in detail the *current* state of the system in order to steer it successfully. One must also have a description of all possible *nearby* states of the system, both under therapeutic pressure and without therapy. Better yet is to have a global picture of *all* possible states of the system, with nonlinear nullcline information, as one would obtain by analyzing the full phase space of the entire system. With this information, one would know *where* to steer the system to get to a desired state, even if one does not know *how* to achieve this (clinically). In current state-of-the-art medical practice, such sophisticated sensor-actuator capability is not yet sufficiently developed as it is in many engineering contexts where adaptive control theory is routinely used. Many similar challenges, and the necessary steps towards their implementation, present themselves in the ecology and pest control communities, and we point to Gould’s article [42] for a nice early overview. More recently, connections between the approaches developed in the past by ecologists and possible future strategies for oncologists have been discussed by Gatenby and collaborators [43]. Other groups [44, 45, 46] have also developed highly mathematical approaches to tumor control from different points of view. Clearly not all of the clinical steps are in place to effectively test and implement many of the strategies that have been explored theoretically. Yet it is still important to continue to develop the kinds of mathematical models and computer simulations that would serve to identify the many possible schemes, parameter ranges, and sensitivities that could be tested via clinical trials that focus on adaptive therapies with the goal of suppression of potential evolution of resistance.

## Footnotes

1 It is important to note that both high-dose, maximum tolerated dose schedules and low-dose, metronomic dose schedules have this cumulative goal of achieving maximum cell-kill over the course of many cycles of treatment.

